# Obesity-related alterations in plasma metabolomics and fecal microbiota in Down syndrome Dp(16)1Yey mice

**DOI:** 10.64898/2026.04.10.717726

**Authors:** Pinku Halder, Mohammed Selloum, Farid Ichou, Loic Lindner, Laura Desnouveaux, François-Xavier Lejeune, Guillaume Pavlovic, Yann Hérault, Marie-Claude Potier, GODS-21 Consortium

## Abstract

**Background/Objectives:** Individuals with Down syndrome (DS) are at increased risk of obesity and metabolic comorbidities, yet the mechanisms underlying these conditions remain unclear. Here we investigated how DS-associated genetic condition interacts with diet and metabolic pathways in the Dp(16)1Yey mouse model of DS.

**Methods:** Untargeted plasma metabolomics was performed in Dp(16)1Yey and control mice, subjected to either control or high-fat diet (HFD). Raw data were processed, and features were annotated. Statistical analyses were conducted in R, and pathway analysis was performed with MetaboAnalyst v5.0. Fecal microbiome was obtained using 16SrRNAseq and analyzed using phyloseq in R.

**Results:** Diet exerted the strongest effect on mice plasma metabolome, followed by sex and genotype. Seventy-five diet-responsive metabolites were enriched in amino acid and nucleotide metabolism. Genotype-driven changes affected 34 metabolites, notably impacting amino acid and taurine–hypotaurine metabolism. Fifty-six sex-associated metabolites highlighted disruptions in aromatic amino acid biosynthesis and pyrimidine metabolism. A significant Diet*Genotype interaction was observed for five metabolites, including a marked reduction in the microbiota-derived metabolite 3-indolepropionic acid (IPA) in Dp(16)1Yey mice on HFD. Both genotype and diet exerted pronounced effects on fecal microbiome with selective depletion of the IPA-producing *Clostridia* in Dp1Yey mice under HFD.

**Conclusion:** Segmental trisomy in Dp(16)1Yey mice modulates the host metabolic response to dietary fat, partly through microbiota-derived metabolites such as IPA. These findings highlight the importance of genotype, diet, and microbiome interactions in shaping metabolic disease risk in DS and point toward microbiota-targeted dietary interventions.

## 1. Introduction

Down syndrome (DS) is the most common genetically derived intellectual disability (ID), resulting from the presence of an extra copy of human chromosome 21 (HSA21), also referred to as trisomy 21 (T21). DS is characterized by a range of core features, including cognitive deficits, craniofacial dysmorphology, hypotonia, and the development of Alzheimer’s disease-like pathology in mid-life. In addition to these hallmark traits, individuals with DS exhibit a spectrum of other abnormalities with varying degrees of penetrance and expressivity, such as congenital heart defects, hearing and vision impairment, leukemia, reduced bone density, and gastrointestinal disorders [1,2]. While intellectual disability remains the most prominent manifestation of DS, significant research has focused on elucidating the effects of T21 on brain function and development [3–5]. However, as the life expectancy of individuals with DS continues to increase [6], a broader range of health challenges has emerged, including metabolic conditions such as obesity, diabetes, and dyslipidemia [7–11]. Notably, body mass index (BMI) and percentage of body fat (%BF) are consistently elevated in individuals with DS relative to age- and sex-matched controls without DS [12,13]. Sex-specific differences are also evident, with females showing disproportionately higher BMI, %BF, and rates of overweight compared to males [14,15]. In addition, individuals with DS have a significantly higher prevalence of Metabolic dysfunction-associated steatotic liver disease (MASLD) a condition strongly associated with obesity and insulin resistance [10]. Although the higher prevalence of obesity, insulin resistance, and diabetes in adolescent and adult DS populations is well documented [8,9,15–18], the underlying mechanisms contributing to impaired metabolic homeostasis in DS remain poorly understood and insufficiently studied.

DS could be considered a metabolic disorder since alterations in metabolites are present in blood and urine from subjects with DS, suggesting that specific disruptions in metabolic pathways may contribute to the pathogenesis of DS. For instance, a Nuclear Magnetic Resonance (NMR) study by Caracausi et al. [19] found that in DS plasma, levels of acetate, acetoacetate, acetone, creatine, formate, glutamine, glycerol, pyruvate, and succinate were significantly elevated, while lysine and tyrosine were reduced. In urine, phenylacetylglycine, trimethylamine-N-oxide (TMAO), and tyrosine were significantly increased, whereas glycine was decreased compared to control subjects [19]. These metabolomic alterations are associated with mitochondrial dysmetabolism. A recent metabolomic study of DS plasma by Liquid Chromatography Mass Spectrometry (LC-MS) analysis highlighted alterations in methylation cycle metabolites, including elevated levels of cystathionine, cysteine, choline, dimethylglycine, S-adenosylhomocysteine (SAH) and S-adenosylmethionine (SAM) [20], although earlier findings reported reduced concentrations of SAH and SAM [21]. These observations further highlight the metabolic dysregulation associated with DS. Such discrepancies underscore the heterogeneous and multifactorial nature of DS, where genetic, epigenetic, and metabolic factors interact to shape disease complexity.

In addition, individuals with DS exhibit impairments in lipid metabolism. Existing studies indicate that children with DS have elevated levels of circulating cholesterol, low-density lipoproteins (LDL), and triglycerides compared to age-matched controls [22–24]. These unfavourable lipid profiles may contribute to the heightened risk of cerebrovascular events [23,25] and the prevalence of overweight and obesity observed in this population [8,11]. Whether these lipid abnormalities stem from the overexpression of specific genes on HSA21 remains an open question. Plasma leptin levels, a key hormonal regulator of energy balance and fat accumulation, show variability in DS, with conflicting reports of both reduced levels in DS foetuses, adolescents and adults compared to controls and elevated levels relative to unaffected siblings [26–29]. Genetic predisposition to leptin resistance may underlie hyperleptinemia observed in DS without hyperinsulinemia [30]. Metabolomics analysis of red blood cells (RBCs) by LC-MS from individuals with DS revealed increased conjugated and deconjugated bile acids, particularly in women compared to control subjects [31], suggesting microbiome alterations [32]. Elevated RBC levels of pyruvate, lactate, and acyl-conjugated fatty acids were also observed in T21 [31], showing an energy metabolism disorder aligning with prior reports of n-6 fatty acid dysregulation in T21 RBCs [33].

DS has been linked to altered mitochondrial respiration, largely attributed to dosage imbalance of HSA21 genes, such as *CBS*, *RCAN1*, *DSCAM*, and *NRIP1*, which impair mitochondrial fusion/fission dynamics and OXPHOS complex function [34–38]. However, these findings are primarily derived from cell culture studies, leaving it unclear whether similar mitochondrial deficits occur in metabolically active tissues, including hepatocytes, myocytes, and adipocytes. A phosphorus magnetic resonance spectroscopy study demonstrated mitochondrial dysfunction in skeletal muscle of adults with DS [39].

The impact of T21 on whole-body metabolism is poorly understood, with most studies focusing on adiposity, food intake, physical activity, and energy expenditure in adolescents and adults with DS [40–44]. Thus, it remains unclear whether metabolic changes are driven by altered gene expression in peripheral tissues due to the extra copy of HSA21 or by lifestyle differences. Recent study in targeted plasma metabolomics analysis in individuals with DS revealed broad metabolic disruption, including impaired lipid and amino acid metabolism and upregulation of the kynurenine pathway of tryptophan catabolism, with interferon (IFN) associated elevations of 5-HIAA, kynurenine, and quinolinic acid [45].

Mouse models carrying segmental duplications of regions orthologous to Hsa21 have been instrumental in dissecting genotype-phenotype correlations in DS [46–49]. Among them, the Ts65Dn model has been extensively used to study neurological and metabolic phenotypes [46]. Studies using this model have shown that high-fat diet (HFD) exposure induces obesity-related biomarkers such as galectin-3 and HSP72, lowers circulating IL-6, and reveals an impaired metabolic-inflammatory axis [50]. Additional reports describe brain insulin resistance, mitochondrial dysfunction, and oxidative stress in Ts65Dn mice, offering mechanistic links to AD risk [51]. HFD-fed Ts65Dn mice also show insulin resistance, adipose tissue inflammation, hepatic pro-fibrotic gene signatures, and disrupted oxidative stress responses [52]. Although these findings underscore the metabolic vulnerability in DS models, the Ts65Dn model includes a non-Hsa21-related segment and does not fully recapitulate the genetic profile of human T21 [53]. In contrast, the Dp(16)1Yey (noted Dp1Yey) mice, encompassing only orthologs of Hsa21 genes exhibited alterations in TCA cycle intermediates and anaplerotic amino acids [54,55]. Similarly, the Dp(16)1Tyb mice, genetically comparable to Dp1Yey, have shown pre-diabetic liver pathology, offering further insight into DS-associated metabolic dysfunction [56]. Additional studies have reported disrupted tryptophan catabolism and monoaminergic neurotransmitter metabolism in the Ts65Dn and Dp1Yey mice [54,55].

Despite the availability of these models, only few studies have systematically assessed the effects of trisomy on baseline metabolic states and adaptive responses to nutritional stress. Notably, there is a lack of detailed metabolomic analyses of the Dp1Yey model, which encompasses the entire HSA21 orthologous region on mouse chromosome 16 and is considered the most genetically faithful DS model. To address this gap, we performed untargeted metabolomics analysis in plasma from Dp1Yey mice subjected to either a normal chow diet (CTL) or HFD. Our study aimed to investigate how segmental trisomy influences systemic metabolic responses under both baseline and obesogenic conditions, with specific attention to sex as a biological variable. By integrating genotype, diet, and sex, this work seeks to provide new insight into early metabolic vulnerabilities in DS and uncover pathways contributing to obesity risk. Finally, we present data on the feces microbiota of a subset of mice to validate changes observed at the metabolomic level.

## 2. Materials and Methods

### 2.1. Experimental animal and dietary intrusion

Male and female Dp1Yey (mutant) mice and their C57BL/6J littermates (WT) were maintained on a standard chow diet (SAFE; D04, Augy, France) for 6 weeks, starting from 4 weeks of age. At 10 weeks of age, the mice were randomly assigned to one of two dietary intervention groups: the CTL group, which continued on the standard chow diet, or the HFD group, which received a high-fat diet (D12451, 45% kcal from fat, Research Diets Inc., New Brunswick, NJ, USA) for a duration of 19 weeks. Each dietary group consisted of a minimum of 7 animals per sex to ensure sufficient statistical power for subsequent analyses.

Animals were housed in groups of four per cage (2 mutants + 2 controls) under controlled environmental conditions: temperature of 21 ± 2 °C, relative humidity of 60 ± 5%, and a 12-hour light-dark cycle. Food and water were provided ad libitum. All animal experiments were conducted in accordance with European Union Directive 2010/63/EU (September 22, 2010) and were approved by the ethics committee of the GODS21 project (approval number: 30859).

### 2.2. Plasma collection for metabolomic

At 29 weeks, following a 4-hour fasting period, mice were anesthetized using isoflurane. Blood samples were obtained via the temporo-mandibular veinpuncture using a needle and immediately transferred into lithium heparin microvette tubes (Sarstedt, Nümbrecht, Germany). To isolate plasma, the collected blood was centrifuged at 2,900g for 15 minutes at 4°C. The resulting plasma samples were aliquoted and stored at −80°C until further analysis.

### 2.3. Chemical reagents for metabolomic

All LC-MS grade reference solvents, acetonitrile (ACN) and methanol (MeOH) were from VWR International (Plainview, NY). LC grade 2-propanol, ammonium carbonate and hydroxide were from Sigma-Aldrich (Saint Quentin Fallavier, France). Stock solutions of stable isotope-labeled mix (Algal amino acid mixture-13C, 15N) were purchased as well from Sigma-Aldrich (Saint Quentin Fallavier, France).

### 2.4. Plasma sample preparation for metabolomic

Eight volumes of frozen acetonitrile (−20 ◦C) containing stable isotope-labeled internal standards (Algal amino acid mixture-13C, 15N; Sigma-Aldrich, Saint Quentin Fallavier, France) at a final concentration of 12.5 µg/mL were added to 50 µL plasma samples and vortexed. After vortexing, samples were sonicated (Branson Ultrasonic BathM3800) for 10 minutes and centrifuged at 10,000 × g for 2 minutes at 4°C. Centrifuged samples were incubated at 4°C for 1 hour for slow protein precipitation and were then centrifuged again at 20,000 × g for 20 minutes at 4°C. 250µl of supernatants were transferred to another series of eppendorf tubes and dried under SpeedVac and stored at −80°C until the LC-MS analysis. Finally, samples were reconstituted in 100 µL of 20/80 (v/v) water/acetonitrile and then centrifuged at 20,000 × g for 10 minutes at 4°C and transferred to vials before LC-MS analyses.

### 2.5. Metabolomic data processing and metabolite identification

LC-MS experiments were performed using HILIC chromatographic column, Sequant ZIC-pHILIC column 5µm, 2.1 × 150 mm at 15◦C (Merck, Darmstadt, Germany). UPLC® Waters Acquity (Waters Corp, Saint-Quentin-en-Yvelines, France) and Q-Exactive mass spectrometer (Thermo Scientific, San Jose, CA) were used and experimental settings were carried out as detailed in Garali et al. [57]. LC-MS raw data files were converted to mzXML format using MSconvert [58]. Peak detection, alignment, correction, and integration were conducted using the XCMS R package with the CentWave algorithm [59,60] and the Workflow4Metabolomics platform [61]. The resulted datasets were log-10 normalized, filtered and cleaned based on quality control (QC) samples [62]. Ion features were annotated based on retention time and mass-to-charge ratio (m/z) using an in-house database [63]. Putative annotations were assigned using public resources, including the Human Metabolome Database (HMDB) [64] and the Kyoto Encyclopedia of Genes and Genomes database, KEGG [65].

After curation, the final dataset consisted of 943 features, with 492 and 451 features detected in ZIC-pHILIC positive and negative ion modes, respectively. From these, 99 metabolites were identified in positive ion mode, and 48 metabolites were identified in negative ion mode. Both datasets were merged for comprehensive analysis of the metabolic profile.

### 2.6. Statistical analyses for metabolomic

All statistical analyses were conducted in R version 4.3.2 (R Development Core Team, 2023). Principal Component Analysis (PCA) was performed on the metabolomics dataset using the ‘prcomp’ function in R, with variables scaled to unit variance. The**factoextra** (v1.0.7) package was used to extract and visualize PCA results. The analysis focused on the first three principal components (PC1–PC3), with PC1 and PC2 capturing the major variance related to diet and sex, and PC3 contributing to genotype-related separation. Individual samples and metabolite variables were projected onto the PCA space and visualized using color and shape codes to indicate diet, genotype, and sex. Multivariate linear regression (MLR) analyses were performed to examine the effects of Diet, Genotype, and Sex on metabolite levels, with each factor adjusted for the others and for body weight as a continuous covariate. To evaluate potential interactions between Diet and Genotype, additional MLR models were considered including Diet × Genotype interaction terms, with Sex as a covariate. MLR models were fitted for each metabolite, the significance of main and interaction effects was assessed using Type II F-tests via the Anova function from the car package (v3.1-2), and p-values were adjusted for multiple comparisons using the Benjamini–Hochberg (BH) method. Metabolites with adjusted p-values < 0.05 were considered statistically significant. Overall patterns of significant metabolite alterations were visualized using heatmaps generated with the pheatmap package (v1.0.12), applying hierarchical clustering based on Euclidean distance and complete linkage.

### 2.7. Pathway analyses of metabolites

Pathway analysis of the significant metabolites was performed using MetaboAnalyst v5.0 (https://genap.metaboanalyst.ca/)[66], where KEGG IDs of metabolites were entered with one compound per row. The enrichment method applied was the Hypergeometric Test, and the topology measure used was Relative-Betweenness Centrality. The ‘Mus musculus (mouse) KEGG’ pathway library was selected for both pathway enrichment and topology analysis. Pathways were ranked based on p-values from the enrichment analysis, with p-values corrected using the Holm-Bonferroni method (pHolm). Pathway impact values were calculated from the topology analysis, and pathways with an impact value > 0.1 were considered the most relevant. Results were visualized as a bubble plot, with bubble size and color representing pathway impact and –log10(p-values), respectively.

### 2.8. Samples collection for metagenomics

Fresh fecal samples were collected from mice at 30 weeks and stored immediately at –80°C until processing.

### 2.9. 16S rRNA Gene Sequencing from fecal samples

Microbial DNA was extracted from fecal samples using the NucleoSpin® DNA Stool kit (Macherey-Nagel, Germany), following the manufacturer’s protocol. DNA quality and concentration were assessed using a NanoDrop spectrophotometer (Thermo Scientific). The extracted DNA samples were sent to the Plateforme d’analyses génomiques at Université Laval (Québec, Canada), where amplification of the V3–V4 hypervariable regions of the 16S rRNA gene was performed using the primers 341F (CCTACGGGNGGCWGCAG) and 806R (GGACTACNVGGGTWTCTAAT). PCR products were then purified, and paired-end sequencing (2×300 bp) was carried out on an Illumina MiSeq platform.

### 2.10. RNAseq data processing

Sequencing data were processed using the FROGS (Find, Rapidly, OTUs with Galaxy Solution version 4.0.1) [67] pipeline implemented on the Galaxy platform (https://usegalaxy.fr/). Reads were quality-filtered, dereplicated, and clustered into Operational Taxonomic Units (OTUs).

Chimera removal and taxonomic assignment were performed using the SILVA reference database (version 138.1). An OTU abundance table was generated for downstream analysis.

### 2.11. Microbiota identification and statistical analyses

Microbial diversity analyses and visualizations were conducted using R (version 4.4.1) and the phyloseq package within the RStudio environment. Alpha diversity was assessed using measures of richness, the Shannon index, and the inverse Simpson index. Beta diversity was calculated based on Bray–Curtis dissimilarity, Jaccard distance, unweighted UniFrac, and weighted UniFrac metrics. Differences between groups were evaluated using PERMANOVA, with p-values < 0.05 considered statistically significant.

## 3. Results

### 3.1. Sex-related alterations in plasma metabolite profile in Dp1Yey and WT mice

To explore the influence of sex on plasma metabolic profiles in Dp1Yey and WT mice, PCA was performed on the 943 detected ion features (see 2.5). Male and female samples showed distinct clustering, primarily along PC2, which accounted for 13.1% of the variance, while PC1 explained 14.9% of the variance (Figure 1A). This separation suggests that sex contributes to variability in the plasma metabolome.

**Figure 1.**
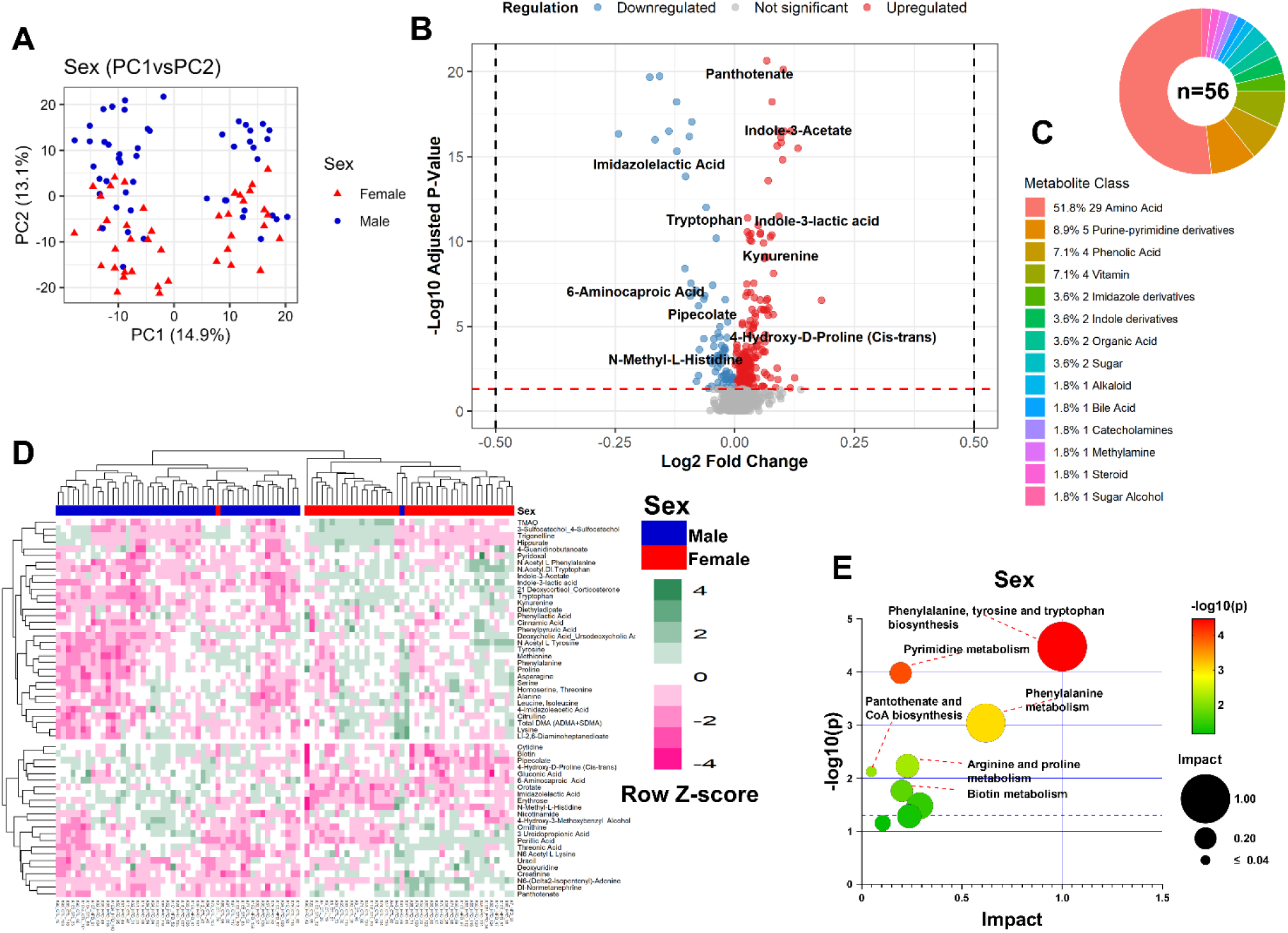
Sex-specific metabolic profile in plasma of Dp1Yey and WT mice. (A) PCA of plasma metabolomic profiles based on all detected metabolites, showing clear separation between Female and Male groups in PC2. (B) Volcano plot showing metabolites significantly altered between sexes using Type II ANOVA F-tests with sex as the main factor, adjusted for diet, genotype and BW (BH adjusted *p* < 0.05). (C) Classification of the 56 significantly altered metabolites into major biochemical pathways, amino acids being the major group (n = 29), followed by purine/pyrimidine derivatives (n = 5), phenolic acids (n = 4), and vitamins (n = 4). (D) Hierarchical Cluster Analysis of significantly altered metabolites reveals sex-dependent sample grouping. Each cell represents the standardized (row Z-score) intensity of a metabolite across samples. Column names indicate mouse IDs. (E) Pathway enrichment analysis of sex-associated metabolites reveals significant perturbation in Phenylalanine, tyrosine, and tryptophan biosynthesis and pyrimidine metabolism (pHolm *p* < 0.05).

Subsequent ANOVA, with sex as the primary factor and adjustments for diet, genotype, and body weight (BW), identified 275 ion features showing significant sex-associated differences. Annotation of these features resulted in 56 identified metabolites (Table S1), of which 45 were elevated and 11 reduced in females relative to males (Figure S1). These sex-related metabolites were visualized in a volcano plot contrasting fold-change and adjusted significance (Figure 1B). These metabolites spanned multiple biochemical classes, with amino acids being the most prominent group (n = 29), followed by purine/pyrimidine derivatives (n = 5), phenolic acids (n = 4), and vitamins (n = 4) (Figure 1C). Hierarchical clustering supported the sex-based distinctions, with a heatmap showing consistent grouping of samples by sex (Figure 1D). Metabolic pathway analysis of the 56 significant metabolites highlighted two key sex-affected pathways: Phenylalanine, tyrosine, and tryptophan biosynthesis (pHolm = 0.0026, impact = 1.0) and pyrimidine metabolism (pHolm = 0.0082, impact = 0.1946) (Figure 1E, Table S2).

These findings underline robust sex-dependent metabolic alteration in mice, emphasizing the need for sex-specific analysis in metabolic studies.

### 3.2. Segmental trisomy induces significant metabolic alterations in Dp1Yey mice

To assess the impact of segmental trisomy on plasma metabolic profiles in Dp1Yey mice, we revisited the PCA of the 943 ion features, this time color-coding individuals by genotype. While PC1 did not distinguish genotypes clearly, separation became more apparent along PC2 and PC3, with PC3 explaining 8.6% of the variance (Figure 2A), indicating that segmental trisomy in Dp1Yey mice results in unique metabolic perturbations. ANOVA with genotype as the primary variable and adjustments for diet, sex, and BW, revealed that 221 out of 943 ion features were significantly altered by the segmental trisomy. Feature annotation resulted in the identification of 34 metabolites (Table S3), with 24 upregulated and 10 downregulated in the Dp1Yey mice compared to WT (Figure S2). A volcano plot visualized these differential metabolites, highlighting the most significantly altered compounds based on fold-change and adjusted p-values (Figure 2B). The metabolites were categorized into distinct biochemical classes, with amino acids comprising the largest group (13 metabolites), followed by acylcarnitines (5 metabolites) and bile acids (4 metabolites) (Figure 2C). Hierarchical clustering confirmed these genotype-specific metabolic differences, with a heatmap (Figure 2D) showing clear genotype-driven clustering patterns.

**Figure 2.**
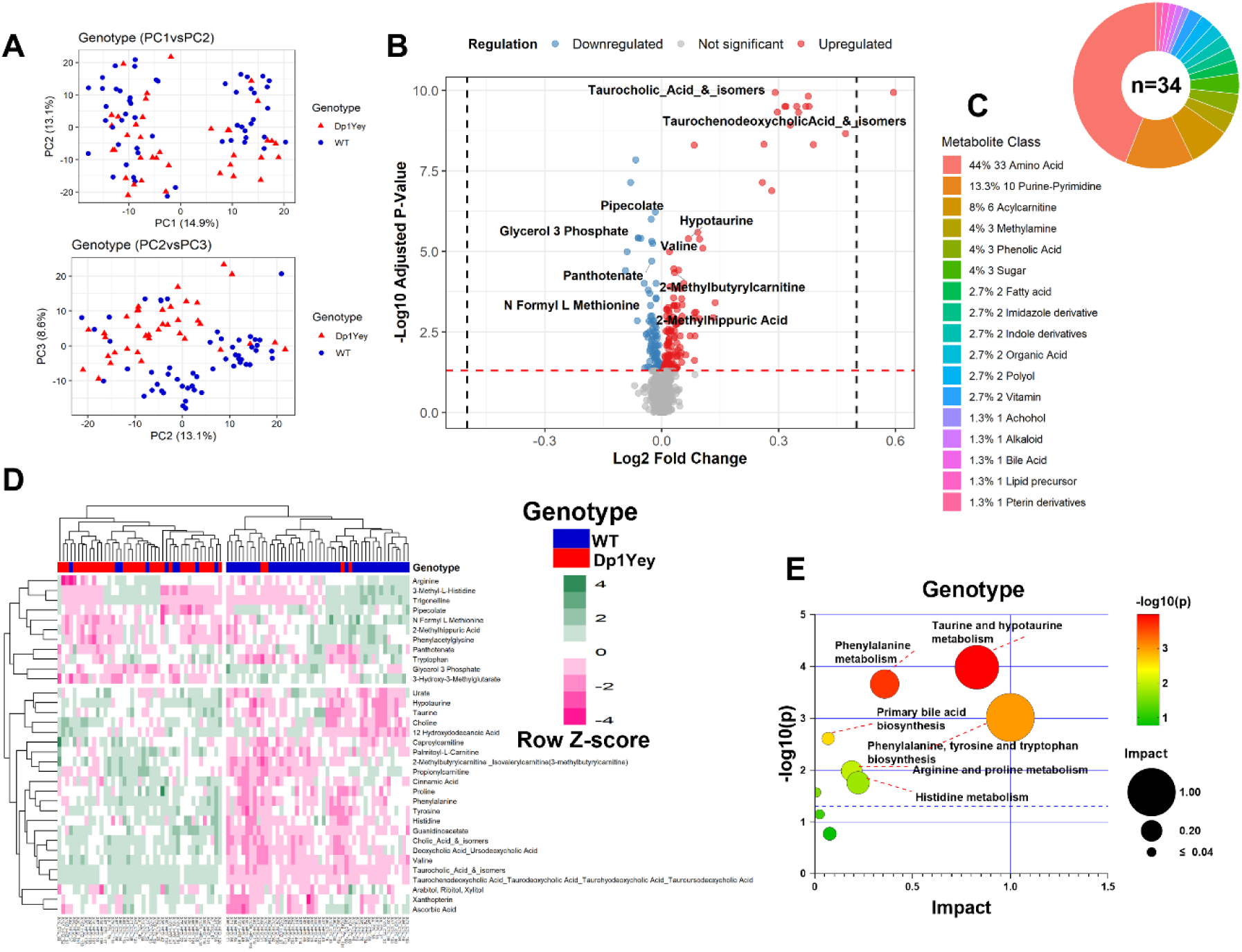
Segmental trisomy induces significant metabolic changes in Dp1Yey mice. (A) PCA plot depicting separation between WT and Dp1Yey groups in PC2 and PC3 not in PC1 and PC2. (B) Volcano plot indicating metabolites significantly altered by genotype using ANCOVA with genotype as the main factor, adjusted for diet, sex and BW (BH adjusted *p* < 0.05). (C) Classification of the 34 significantly altered metabolites into major biochemical pathways, with amino acids comprising the largest group (13 metabolites), followed by acylcarnitines (5 metabolites) and bile acids (4 metabolites). (D) Hierarchical Cluster Analysis of significantly altered metabolites reveals genotype-dependent sample grouping. Each cell represents the standardized (row Z-score) intensity of a metabolite across samples. Column names indicate mouse IDs. (E) Pathway enrichment analysis of genotype-associated metabolites indicates involvement of taurine and hypotaurine and phenylalanine metabolism (pHolm *p* < 0.05).

Metabolic pathway analysis of the 34 significant metabolites uncovered two key pathways disrupted in Dp1Yey mice compared to WT controls. Taurine and hypotaurine metabolism (pHolm = 0.0083, impact = 0.829) showed the greatest perturbation, followed by phenylalanine metabolism (pHolm = 0.017, impact = 0.357) (Figure 2E, Table S2).

Together, these results demonstrate that segmental trisomy in Dp1Yey mice induces significant metabolic alterations independently of sex, diet and BMI, particularly affecting amino acid and taurine and hypotaurine metabolism.

### 3.3. High-Fat Diet induces broad changes in plasma metabolite profiles in Dp1Yey and WT mice

To investigate the impact of a HFD on plasma metabolic homeostasis in Dp1Yey and WT mice, we revisited the previously performed PCA of 943 detected ion features, this time visualizing samples by diet group. The analysis revealed a clear separation between CTL and HFD groups along the PC1 (Figure 3A). This separation reflected distinct dietary-dependent metabolic profiles, highlighting the robust metabolic shift induced by HFD exposure.

**Figure 3.**
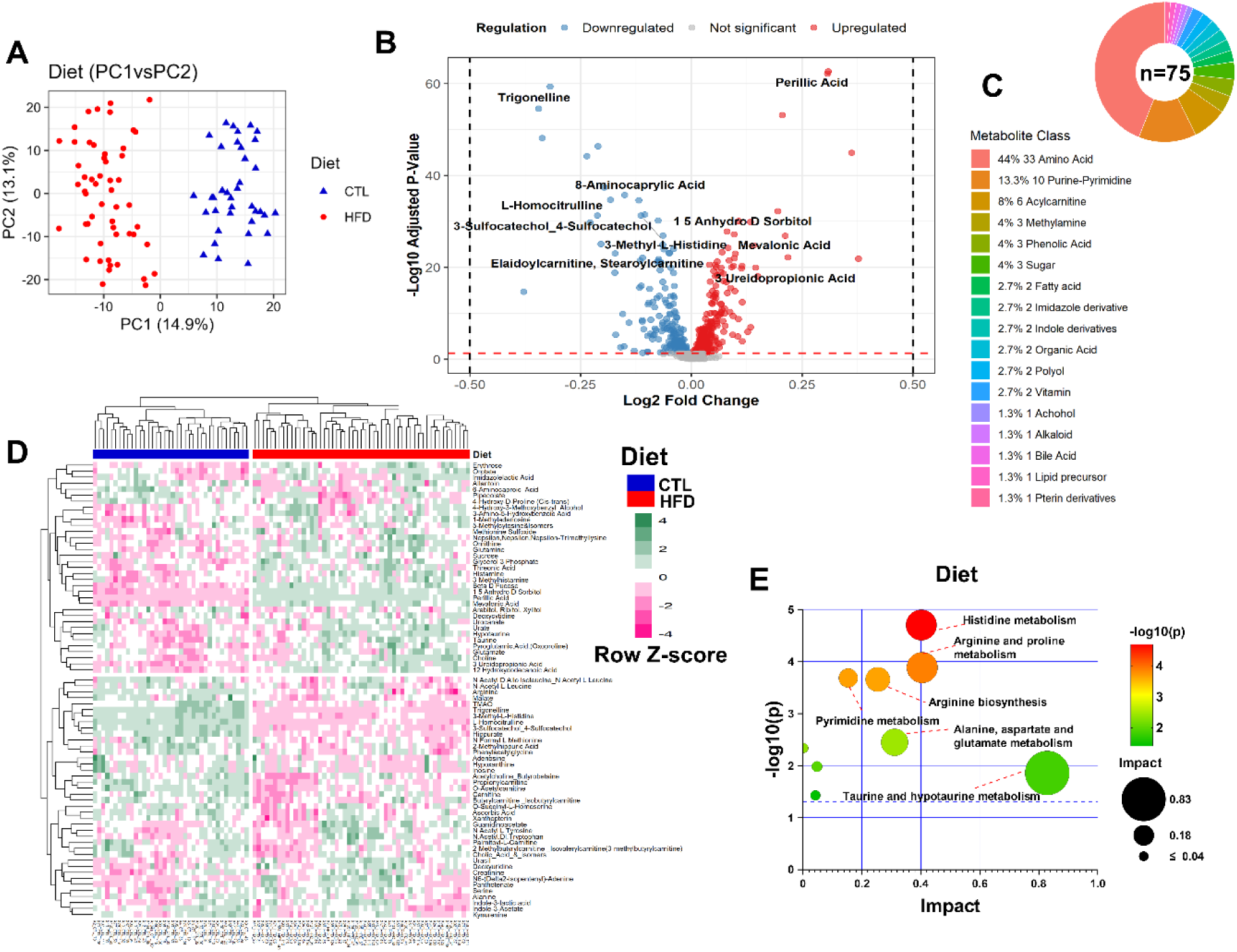
High-fat diet induces widespread metabolic remodelling in Dp1Yey and WT mice. (A) PCA of plasma metabolomic profiles based on all detected metabolites, showing clear separation between HFD and CTL groups. (B) Volcano plot highlighting metabolites significantly altered in HFD versus CTL groups using ANOVA with diet as the main factor, adjusted for genotype and sex, and corrected for multiple testing using the BH method (adjusted *p* < 0.05). (C) Classification of the 75 significantly altered metabolites into major biochemical categories, with amino acids being the most represented class. (D) Hierarchical clustering analysis of significantly altered metabolites reveals diet-dependent sample grouping. Each cell represents the standardized (row Z-score) intensity of a metabolite across samples. Column names indicate mouse IDs. (E) Pathway enrichment analysis based on the 75 significant metabolites, identifies histidine metabolism, arginine and proline metabolism, pyrimidine metabolism, and arginine biosynthesis as significantly impacted pathways (pHolm *p* < 0.05).

To assess the global impact of diet on metabolite abundance, we performed ANOVA, using diet as the main factor while adjusting for sex and genotype. A total of 467 out of 943 ion features were significantly altered under the HFD condition. Subsequent annotation resulted in the identification of 75 unique metabolites (Table S4), of which 40 were upregulated and 35 were downregulated in the HFD group compared to CTL (Figure S3). A volcano plot visualized these metabolite shifts, emphasizing the most significantly affected compounds based on fold-change and adjusted p-values (Figure 3B).

The 75 significantly altered metabolites were further classified into biochemical categories, with amino acids comprising the largest group (n = 33), followed by purines/pyrimidines (n = 10) and acylcarnitines (n = 6), among others (Figure 3C). These significantly altered metabolites were visualized using hierarchical clustering, which showed distinct grouping of samples by dietary condition (Figure 3D), consistent with the PCA results and further confirming the global impact of HFD on the plasma metabolome of mice.

Pathway enrichment analysis of the 75 significant metabolites identified four metabolic pathways that were significantly impacted by the HFD compared to the CTL group: histidine metabolism (pHolm=0.0016, impact = 0.402), arginine and proline metabolism (pHolm=0.010, impact = 0.405), pyrimidine metabolism (pHolm=0.016, impact = 0.154), and arginine biosynthesis (pHolm=0.017, impact = 0.254) (Figure 3E, Table S2).

Collectively, these findings demonstrate that HFD exposure induces significant remodelling of the plasma metabolome in Dp1Yey mice, with specific alterations in amino acid and nucleotide metabolic pathways, indicating a broad reprogramming of metabolic processes in response to the diet.

### 3.4. Overlapping Metabolic Signatures Across Diet, Genotype, and Sex Reveal Specific Pathway Associations

To identify shared metabolic signatures, we overlapped the lists of significantly altered metabolites between diet, genotype, and sex comparisons, focusing on those common to two experimental variables only. The largest overlaps were observed between diet and sex, with 24 shared metabolites, followed by diet and genotype (19 metabolites), and sex and genotype (6 metabolites) (Figure 4A). These overlaps highlight the complex interactions between these variables in shaping metabolic profiles in Dp1Yey mice.

**Figure 4.**
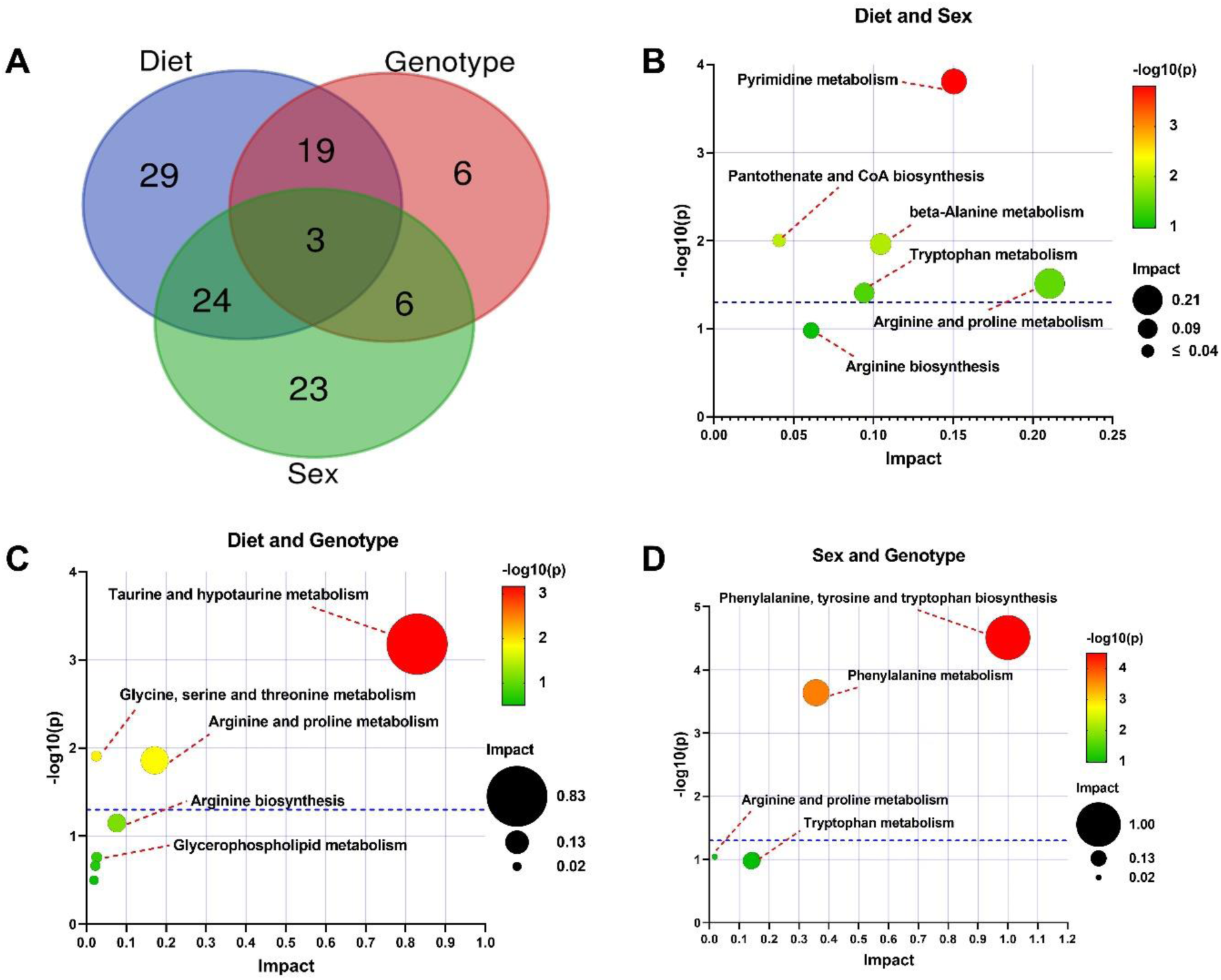
Overlapping metabolic signatures across diet, genotype, and sex reveal pathway-specific associations. (A) Venn diagram illustrating overlap of significantly altered metabolites among diet, genotype, and sex comparisons. (B) Pathway enrichment analysis of 24 metabolites shared between diet and sex reveals significant enrichment of pyrimidine metabolism (pHolm = 0.012). (C) Nineteen metabolites overlapping between diet and genotype show high impact in taurine and hypotaurine metabolism (impact = 0.829), with marginal significance (pHolm = 0.053). (D) Six metabolites shared between sex and genotype map to phenylalanine, tyrosine, and tryptophan biosynthesis (pHolm = 0.0025, impact = 1.0) and phenylalanine metabolism (pHolm = 0.018, impact = 0.357). Pathway analysis was performed using MetaboAnalyst v5.0.

Pathway analysis of the overlapping metabolites revealed distinct associations with key metabolic pathways. Metabolites shared between diet and sex were significantly associated with pyrimidine metabolism (pHolm = 0.012, impact = 0.1504) (Figure 4B, Table S5), underscoring the role of pyrimidine derivatives in the metabolic shifts linked to both dietary and sex-related factors. For the metabolites shared between diet and genotype, taurine and hypotaurine metabolism exhibited the high impact (impact = 0.829), though the association was marginally significant after p-value adjustment (pHolm = 0.053) (Figure 4C, Table S5). This suggests that taurine and hypotaurine metabolism might be influenced by both dietary and genetic factors; however, further investigation is needed to validate this observation. Finally, the metabolites shared between sex and genotype were strongly linked to phenylalanine, tyrosine, and tryptophan biosynthesis (pHolm = 0.0025, impact = 1.0) (Figure 4D, Table S5), which showed the highest impact among the overlapping pathways. Additionally, phenylalanine metabolism was also significantly associated (pHolm = 0.018, impact = 0.357), highlighting the main influence of both sex and genotype on the metabolism of aromatic amino acids.

Together, these results reveal that overlapping metabolic signatures across diet, genotype, and sex lead to specific pathway alterations, further elucidating the complex and multifactorial nature of metabolic regulation in Dp1Yey mice.

### 3.5. Synergistic effect of diet and genotype on metabolite levels reveals associations with microbiota in the Dp1Yey mouse model

To evaluate how plasma metabolite levels are altered in the Dp1Yey mouse model under a high-fat diet (HFD) challenge relative to WT, an ANOVA was performed using the DietxGenotype interaction term, with adjustment for sex. This analysis revealed a significant synergistic effect of diet and genotype on five metabolites: indole-3-lactic acid, 4-guanidinobutanoate, *N*-caffeoylputrescine, 3-indolepropionic acid (IPA), and L-serine (Figure 5A–B, Table S6). Indole-3-lactic acid and IPA, both microbiota-derived products of tryptophan metabolism, staid significant, underscoring the influence of gut microbiota in shaping host metabolic responses. Specifically, IPA levels were significantly reduced in Dp1Yey mice under HFD compared to WT counterparts particularly in males (Figure S4A), reflecting genotype-dependent susceptibility to diet-induced metabolic perturbations. To further explore whether IPA concentrations are linked to host phenotypic traits, we performed sex-stratified Spearman correlation analyses between body weight and plasma IPA levels. This revealed significant inverse correlations in Dp1Yey males (ρ = −0.678, *p* = 0.0007), WT females (ρ = −0.576, *p* = 0.0026), and WT males (ρ = −0.479, *p* = 0.0099), while no significant correlation was observed in Dp1Yey females (ρ = −0.234, *p* = 0.37) (Figure S4B). These results suggest that sex-specific metabolic regulation may modulate the association between host adiposity and microbiota-derived metabolites such as IPA in a genotype-dependent manner.

**Figure 5.**
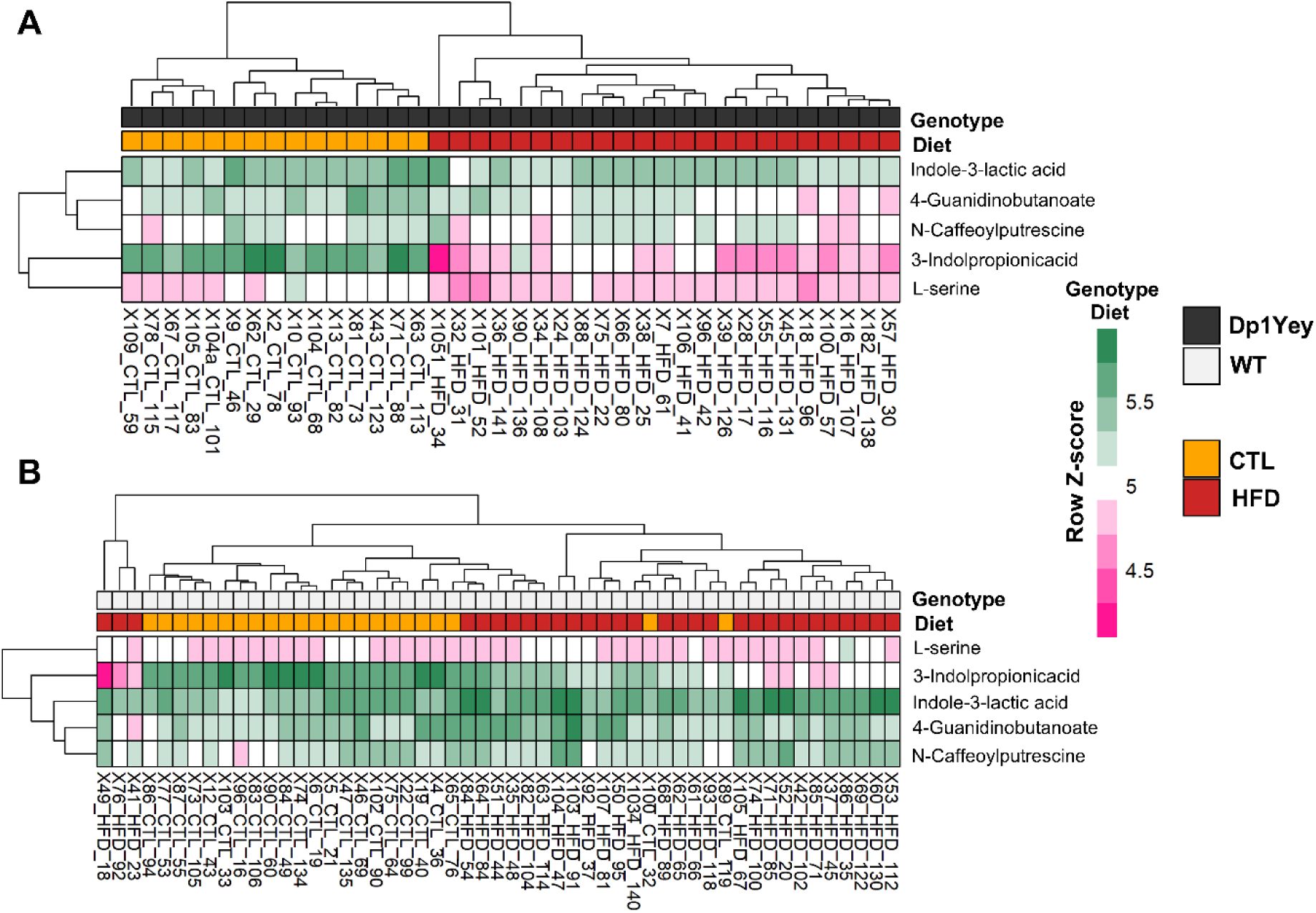
Heatmaps of metabolites significantly altered by Diet × Genotype interaction in Dp1Yey and WT mice. (A) Heatmap showing plasma metabolite profiles in Dp1Yey mice under CTL and HFD conditions. (B) Corresponding heatmap for WT mice under CTL and HFD conditions. Metabolites were identified through F-tests including the Diet×Genotype interaction term, adjusted for sex. P-values were corrected for multiple comparisons using the BH method, with a significance threshold of adjusted *p* < 0.05. The heatmaps demonstrate distinct diet-dependent metabolic responses in Dp1Yey compared to WT mice, highlighting interaction-specific metabolic signatures. Each cell represents the standardized (row Z-score) intensity of a metabolite across samples. Column names indicate mouse IDs.

### 3.6. Combined effects of genotype and diet modulate gut microbial diversity and taxonomic composition in the Dp1Yey mouse model

To investigate the impact of genotype and diet on gut microbial communities in Dp1Yey mice, we quantified overall diversity, β-diversity, and specific taxonomic shifts using 16S rRNA gene profiling of fecal samples. Analysis of OTU numbers (Figure 6A) revealed a significant decrease in microbial richness in both Dp1Yey and WT mice exposed to a HFD compared to their CTL counterparts. These results indicate that both the duplication in the Dp1Yey and dietary intervention synergistically impact on gut microbial diversity.

**Figure 6.**
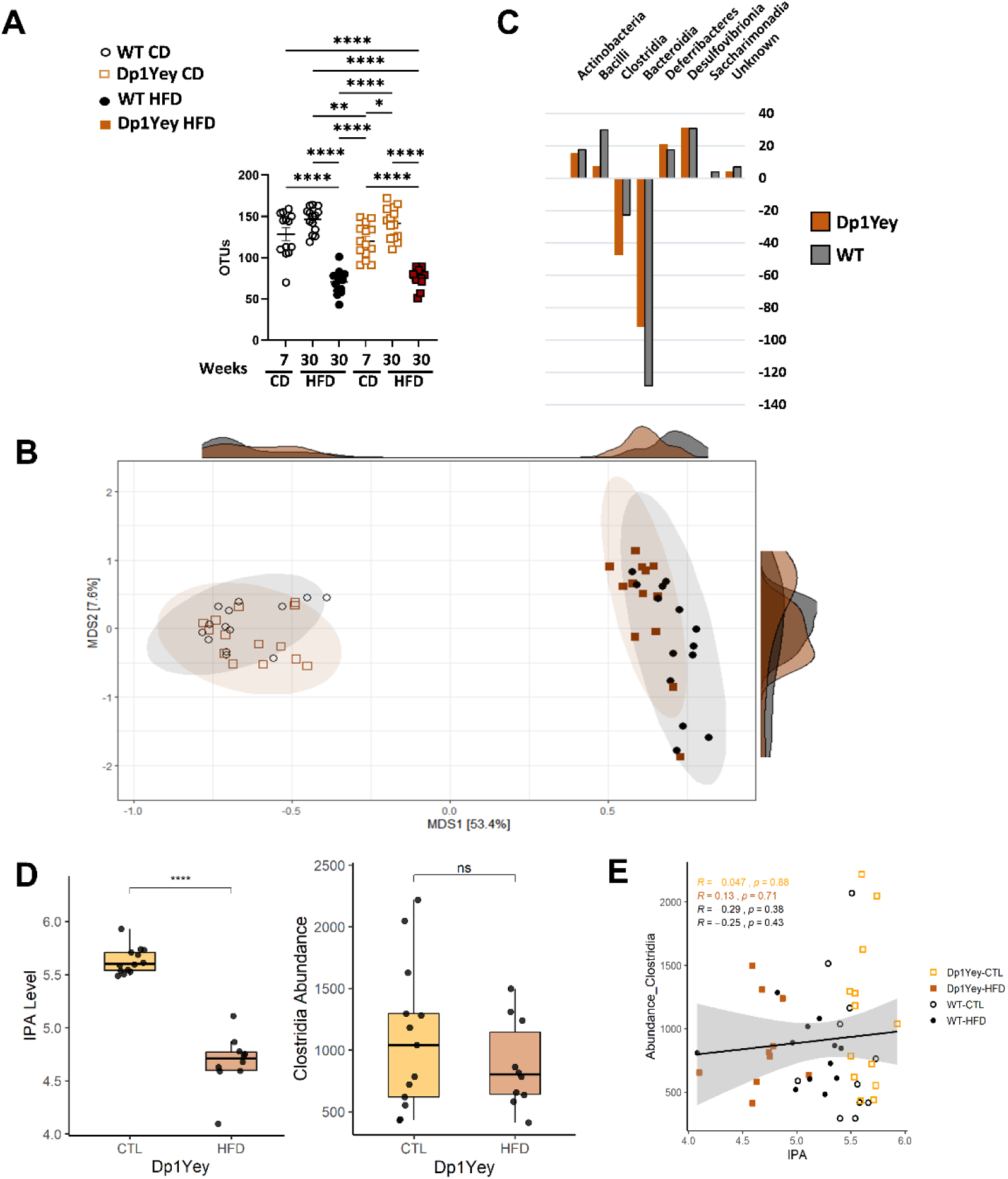
Impact of Genotype and Diet on Gut Microbial Diversity and Composition. **(A)** Number of observed OTUs: Scatter plot showing OTU richness in wild-type (WT) and Dp1Yey mice at 7 and 30 weeks, maintained on CTL or HFD. **(B)** Jaccard index-based multidimensional scaling (MDS) plot visualizing differences in microbiota composition between groups, colored by genotype and diet, with associated density ridges reflecting sample distribution. **(C)** Z-score analysis of bacterial taxa abundance changes: Bar plot comparing Z-score normalized changes in major bacterial taxa between Dp1Yey and WT groups, highlighting a pronounced depletion of Clostridia in Dp1Yey. Positive Z-scores indicate enrichment, while negative values indicate taxa depletion relative to controls. **(D)** Box plots depicting plasma IPA concentrations and fecal Clostridia abundance in Dp1Yey mice under CTL and HFD conditions. **(E)** Scatter plot showing the relationship between Dp1Yey plasma IPA levels and Clostridia abundance in fecal samples. Regression lines with 95% confidence intervals are shown for each group. Spearman correlation coefficients (R) and the corresponding p values are displayed for each group. Statistical significance between groups is indicated by asterisks (*p* > 0.05 = ns*; p* ≤ 0.05 = *; *p* ≤ 0.01 = **; *p* ≤ 0.001 = ***; p < 0.0001 = ****).

Jaccard index-based multidimensional scaling (MDS) (Figure 6B) demonstrated clear segregation of gut microbial communities according to genotype and dietary regimen. Clear group separation along the primary axis (MSD1, 53.4%) demonstrates that Dp1Yey mice and WT controls, subject to varying diets, harbor distinct microbial communities. The clustering pattern suggests that both genetic background and dietary composition exert pronounced effects on gut microbiome structure. Density ridges above and to the side of the MDS plot summarize sample distributions, further supporting the segregation between groups. The observed clustering pattern underscores strong β-diversity differences, contributing to distinct compositional profiles among the groups.

Z-score analysis of taxonomic changes with HFD highlighted a substantial and selective reduction in *Clostridia* abundance in Dp1Yey mice (Figure 6C), markedly exceeding the reduction observed in WT controls. This finding suggests that the Dp1Yey genotype confers heightened susceptibility to *Clostridia* depletion under HFD conditions. Given the critical roles of *Clostridia* in gut health and metabolic regulation, their loss may have functional implications for host physiology in Dp1Yey mice. In contrast, for Bacteroidia, Z-score analysis reveals a moderate decrease in WT mice as compared to Dp1Yey mice (Figure 6C). This comparative shift indicates that the Dp1Yey genotype may partially preserve Bacteroidia populations, potentially as a genotype-specific response to dietary intervention.

In summary, these results reveal that genotype and diet synergistically shape gut microbial diversity and composition in Dp1Yey mice. Notably, *Clostridia* are markedly depleted while Bacteroidia are relatively preserved in the Dp1Yey genotype, highlighting distinct, taxon-specific microbiota responses. These findings underscore the relevance of host genetics and dietary influences in modulating the gut microbiome, with potential implications for metabolic phenotypes in Down syndrome mice models.

### 3.7. Association between plasma IPA levels and gut Clostridia abundance in relation to genotype and diet in the Dp1Yey mouse model

To elucidate the relationship between host metabolites and gut microbiota in the Dp1Yey mouse model, we performed a correlation analysis on a subset of Dp1Yey and CTL mice integrating plasma IPA levels from untargeted metabolomics with *Clostridia* taxon abundance derived from 16S rRNA fecal microbiota profiling at Week 30. Consistent with our overall findings, Dp1Yey mice exposed to a HFD exhibited significantly reduced plasma IPA levels relative to the CTL group (Figure 6D, p < 0.0001), while *Clostridia* abundance in fecal samples showed a downward trend that did not reach statistical significance (p>0.05).

Correlation analysis at 30 weeks revealed that coefficients across Dp1Yey-CTL (R = − 0.047, p = 0.88), Dp1Yey-HFD (R = 0.13, p = 0.71), WT-CTL (R = −0.29, p = 0.38), and WT-HFD (R = −0.25, p = 0.43) all indicated weak, non-significant associations (Figure 6E). Thus, these analyses for each group demonstrate no significant linear relationship between plasma IPA and *Clostridia* levels (all p > 0.05).

Collectively, our results indicate that while Dp1Yey mice exhibit genotype and diet-dependent reductions in both circulating IPA and fecal *Clostridia* abundance, no statistically significant direct correlation was observed. The integrated approach underscores the complexity of host-microbiota metabolic cross talk, suggesting that the modulation of IPA and *Clostridia* may occur via another gut microbiota, or independent or multifactorial mechanisms in this Down syndrome mouse model.

## 4. Discussion

In this study, we performed an untargeted plasma metabolomic analysis in the Dp1Yey mouse model of DS and WT controls, exploring the independent and interactive effects of diet, sex, and genotype. We provide compelling evidence that segmental trisomy (genotype), dietary context, and biological sex independently and interactively contribute to the metabolic landscape of DS in these mice. Using the LC-MS untargeted metabolomics, we profiled 943 plasma ion features across multiple experimental variables, revealing distinct and biologically consistent metabolic signatures. Our multidimensional analyses, encompassing PCA, ANOVA, and multivariate regression models, reveal that these variables not only act independently but also interact synergistically to shape metabolomic outcomes, underscoring the necessity of integrative analysis in genetically complex models of DS.

### 4.1. Diet as the major determinant of metabolic variation

HFD induced the most profound metabolic changes in the plasma metabolomic profile of mice, with significant differences observed in over 467 ion features and 75 annotated metabolites, affecting diverse biochemical pathways including amino acid and nucleotide metabolism. Recent metabolomics studies have reinforced the substantial impact of HFD and related obesogenic diets. Vieira et al. [68] reported in C57BL/6J mice under HFD widespread alterations in nicotinamide, branched-chain amino acid, and cofactor metabolism, with metabolites such as hippuric acid and short chain carnitine (carnitine C4) also decreased in our HFD-fed Dp1Yey mice, supporting the reproducibility of diet-associated metabolic signatures. Similarly, Aggarwal et al. [69] identified HFD-induced disruptions in arginine biosynthesis, pyrimidine metabolism, alanine, aspartate, glutamate metabolism, and the citrate cycle pathways, also perturbed in our HFD-fed Dp1Yey cohort, highlighting conserved metabolic responses to dietary excess. Our data thus reinforce that dietary interventions exert robust effects on metabolic phenotypes, detectable even after adjustment for genotype in a chromosomal aneuploidy model.

### 4.2. Sex-dependent metabolic signatures in mice

Sex was the second most influential variable, with significant differences observed in over 275 ion features and 56 annotated metabolites. Many of these were amino acids, particularly within the phenylalanine, tyrosine, and tryptophan biosynthesis pathways. These aromatic amino acids serve as precursors for monoaminergic neurotransmitters, and their dysregulation may underlie sex-dependent differences in behaviour, immunity, and neurochemistry [70–72]. The enrichment of these pathways supports recent findings that hormonal and hepatic enzyme profiles differ between male and female mice and contribute to metabolic dimorphism [73,74]. Several studies [75–77] describe this sexual dimorphism, especially in obesity. These studies [77–79] emphasize the role of sex hormones in modulating lipid metabolism, the levels of high-density lipoprotein, low-density lipoprotein and triglycerides. Changes in lipid and lipoprotein with body fat distribution, particularly visceral fat, and other still unclear parameters contribute to the differences in cardiometabolic risk between men and women [80]. Our results highlight the need to account for sex as a biological variable in metabolomic studies, especially in models of neurodevelopmental disorders such as DS.

### 4.3. Specific effects of the segmental duplication in Dp1Yey mice

Although the direct effect of the Dp1Yey genotype on plasma metabolite profiles was modest in comparison to diet and sex, we identified 34 annotated metabolites that differed significantly between genotypes. These included acylcarnitines, bile acids, and amino acids, reflecting known alterations in mitochondrial β-oxidation and hepatic function in DS [52].

Among these, choline was significantly elevated in Dp1Yey plasma, consistent with previous reports of reduced folate availability and altered one-carbon metabolism in individuals with DS [20,81]. Disruption of the choline–folate axis has been implicated in both neurodevelopmental and systemic metabolic dysregulation in DS. Furthermore, several acylcarnitines, including valeryl-isovaleryl carnitine (C5), propionyl carnitine (C3), caproyl carnitine (C6), and palmitoyl L-carnitine (C16), were significantly upregulated in Dp1Yey mice. These metabolites, central to mitochondrial fatty acid β-oxidation, support evidence of impaired mitochondrial function in DS, consistent with findings from human plasma metabolomics studies [81,82].

Notably, while Antonaros et al. [83] and Caracausi et al. [19] reported decreased plasma concentrations of tyrosine, histidine, and threonine, along with elevations in energy-related metabolites such as acetate, acetoacetate, creatine, formate, glycerol, pyruvate, and succinate in individuals with DS, we observed increased tyrosine and histidine and no such elevations in energy-related metabolites in Dp1Yey plasma. These discrepancies likely reflect species-specific metabolic regulation or gut microbial contributions.

Another recent human plasma LC-MS study by Colak et al. [82] identified significant downregulation of vitamin C, taurolithocholic acid, pantothenic acid, prostaglandins, cholic acid, and short-chain acylcarnitines (e.g., CAR 10:0, CAR 10:1), alongside upregulation of tauroursodeoxycholic acid, thymidine, serine, nervonic acid, hypoxanthine, and long-chain carnitines (e.g., CAR 16:1, CAR 14:1, CAR 12:0). Our findings partially mirror these alterations, showing increased levels of cholic acid and its isomers, vitamin C, tauroursodeoxycholic acid, and several carnitine derivatives in Dp1Yey plasma, as well as decreased pantothenic acid and arginine. These overlapping and divergent results highlight both conserved and model-specific metabolic features associated with trisomy 21, particularly in bile acid metabolism, mitochondrial fatty acid transport, and vitamin/cofactor homeostasis. Consistent with our findings, [84] reported elevated levels of bile acids, including taurochenodeoxycholic acid and tauroursodeoxycholic acid, in trisomy 21 iPSCs compared to isogenic controls using untargeted metabolomics by LC-MS, suggesting that bile acid dysregulation may be an early, genotype-driven metabolic hallmark of trisomy 21.

Importantly, while the Dp1Yey genotype alone induced limited metabolic alterations, its interaction with HFD amplified selective effects, suggesting a synergistic susceptibility to metabolic stress.

### 4.4. Shared and distinct pathway associations across variables

Intersection analysis revealed distinct metabolic signatures shared across variable pairs. Diet and sex shared the largest set of metabolites (n = 24), with significant enrichment in pyrimidine metabolism, an essential pathway in nucleotide synthesis and energy metabolism. Diet and genotype shared 19 metabolites, prominently affecting taurine and hypotaurine metabolism, again pointing to shared metabolic stress mechanisms. The smallest overlap occurred between sex and genotype (n = 6), but was notably enriched in aromatic amino acid metabolism, emphasizing its relevance in DS and its modulation by sex [55,70].

### 4.5. Diet and Genotype Interactions Reveal Microbiota-Linked Signatures

Importantly, interaction modelling of diet and genotype revealed five metabolites with significant Diet*Genotype interactions, all altered specifically in Dp1Yey mice under HFD. Among them, indole-3-lactic acid and 3-indolepropionic acid (IPA), both microbial metabolites of tryptophan, were altered suggesting an interaction between host genetics and the gut microbiome in modulating systemic metabolism. The role of the microbiota in shaping DS-related phenotypes is increasingly appreciated, including effects on brain development, inflammation, immune and metabolic regulation [85–88] . One of the most noteworthy findings from our analysis was the significant decrease in the levels of IPA in the Dp1Yey group following HFD exposure. IPA, a tryptophan-derived metabolite produced by gut microbiota, has been recognized for its potential protective effects against metabolic diseases, including Type 2 diabetes mellitus (T2DM), insulin resistance and heart failure [89–91]. Furthermore, IPA has been linked to modulation of the gut-brain axis, where it influences energy homeostasis and adiposity through mechanisms such as reducing inflammation and promoting metabolic flexibility [92]. The reduced levels of IPA observed in our study, specifically in the Dp1Yey group, may suggest a compromised microbiota-driven metabolic pathway in these mice under HFD challenge. Importantly, our sex-stratified Spearman correlation analyses revealed significant inverse relationships between body weight and plasma IPA levels in Dp1Yey males, WT females, and WT males, but not in Dp1Yey females. These findings suggest that sex-specific metabolic regulation modulates the interplay between host adiposity and microbiota-derived metabolites such as IPA in a genotype-dependent manner. Studies have shown that lower levels of IPA are associated with increased risk of insulin resistance and metabolic dysfunction [93,94]. Moreover, a recent study reported that short-term HFD exposure leads to decreased IPA levels in the mouse brain, accompanied by changes in metabolites linked to oxidative stress and amino acid metabolism [95]. This association raises the intriguing possibility that the altered microbiota in Dp1Yey mice, in combination with HFD exposure, could contribute to the observed metabolic dysregulation. Interestingly, while the Dp1Yey group exhibited a significant decrease in IPA levels, no such effect was observed in the WT group, further supporting the idea that the segmental trisomy in Dp1Yey mice exacerbates the metabolic perturbations induced by HFD. This genotype-dependent response highlights the genetic predisposition of Dp1Yey mice to altered microbiota-mediated metabolic processes, particularly in the context of dietary interventions. Notably, this genotype dependent effect was more pronounced in male mice, suggesting a sex-specific susceptibility to these metabolic alterations.

The reduced levels of IPA and its association with microbiota in Dp1Yey mice provide a compelling area for future investigation. The microbiota-gut-brain axis and its interaction with metabolic processes could offer insights into the pathophysiology of T2DM and related metabolic diseases. Furthermore, the potential role of microbiota-derived metabolites, such as IPA, in modulating insulin sensitivity and glucose homeostasis warrants further exploration.

Altogether, this integrative metabolomic analysis emphasizes the importance of multidimensional experimental design when evaluating metabolic outcomes. The observed hierarchies—where diet > sex > genotype—underscore the importance of considering modifiable factors, such as nutrition, alongside fixed genetic effects. Moreover, the identification of microbiota-linked metabolites as targets of Diet×Genotype synergy opens promising avenues for gut microbiome modulation as a potential intervention strategy in Down syndrome and related metabolic disorders.

### 4.6. Gut microbiota alterations reflect genotype and diet-dependent metabolic remodeling in Dp1Yey mice

To further interpret the systemic metabolic changes observed in plasma metabolomics, we characterized gut microbial diversity and taxonomic composition in a subset of Dp1Yey and WT mice. Consistent with previous reports demonstrating that HFD profoundly alters intestinal ecology [96,97], our data revealed that both genotype and dietary context jointly shape gut microbial diversity. Specifically, microbial richness was reduced in both WT and Dp1Yey mice upon HFD exposure, indicating that the obesogenic diet imposes a dominant pressure on gut ecosystem complexity. However, multidimensional scaling analyses demonstrated distinct clustering of microbial communities according to genotype and diet, suggesting that segmental trisomy modifies the host’s microbiota response to nutritional stress.

Among the most striking findings was a selective depletion of *Clostridia* in Dp1Yey mice, which was substantially greater than that observed in WT controls under HFD. *Clostridia* are key butyrate-producing bacteria involved in maintaining intestinal barrier integrity, regulating immune tone, and generating tryptophan-derived metabolites including IPA [98–101]. Their loss in Dp1Yey mice may therefore contribute to both local and systemic metabolic perturbations. In contrast, Bacteroidia populations were relatively preserved in Dp1Yey mice, indicating that the effects of trisomy on microbial taxa are selective rather than global. Such taxon-specific responses highlight how genotype influences microbial resilience under metabolic stress.

Despite parallel reductions in both circulating IPA and fecal *Clostridia* abundance, no statistically significant direct correlation between these measures was observed. This lack of a simple linear relationship underscores the complexity of host–microbiota metabolic cross-talk in DS, where genotype-dependent host factors such as altered bile acid metabolism, immune regulation, and intestinal motility may indirectly modulate microbial metabolite production and absorption. Similar uncoupling between microbial abundance and metabolite levels has been observed in other metabolic and neurodevelopmental disorders [102,103], supporting the idea that multiple, interdependent mechanisms govern host–microbiota interactions in DS.

Collectively, these findings suggest that segmental trisomy amplifies the impact of dietary stress on gut microbial communities, leading to selective depletion of beneficial taxa such as *Clostridia* and concomitant reduction of protective microbial metabolites such as IPA. This integrative analysis thus provides a mechanistic link between genotype-dependent microbiota remodeling and metabolic vulnerability in the Dp1Yey model. Future studies combining metagenomic and metabolomic profiling at higher resolution will be instrumental in delineating causal pathways linking microbial dysbiosis, host metabolism, and obesity risk in Down syndrome.

## 5. Conclusion

In conclusion, our integrative metabolomic and microbiota analyses of the Dp1Yey mouse model demonstrate that diet exerts the dominant influence on systemic metabolism, followed by sex and segmental trisomy. These interacting factors collectively remodel key metabolic pathways, particularly those involving amino acid, lipid, and nucleotide metabolism. Among the most notable findings was the genotype-dependent depletion of IPA in Dp1Yey mice under HFD conditions, accompanied by a parallel reduction in fecal *Clostridia* abundance. Given the established role of IPA as a microbiota-derived metabolite with anti-inflammatory, antioxidant, and insulin-sensitizing properties, its decline suggests impaired gut–host metabolic crosstalk, potentially predisposing individuals to insulin resistance, type 2 diabetes, and obesity-related comorbidities (Figure 7). By coupling untargeted metabolomics with targeted microbial community profiling, our results highlight the importance of considering microbiota-host interactions when evaluating metabolic risk in genetically predisposed populations such as individuals with DS. Specifically, the results point to dietary interventions targeting microbial tryptophan metabolism could offer a promising avenue for mitigating metabolic dysfunction in DS models. Future studies integrating metagenomic, metabolomic, and functional assays will be essential to delineate causal pathways and identify microbial or dietary interventions capable of restoring metabolic resilience in DS contexts.

**Figure 7.**
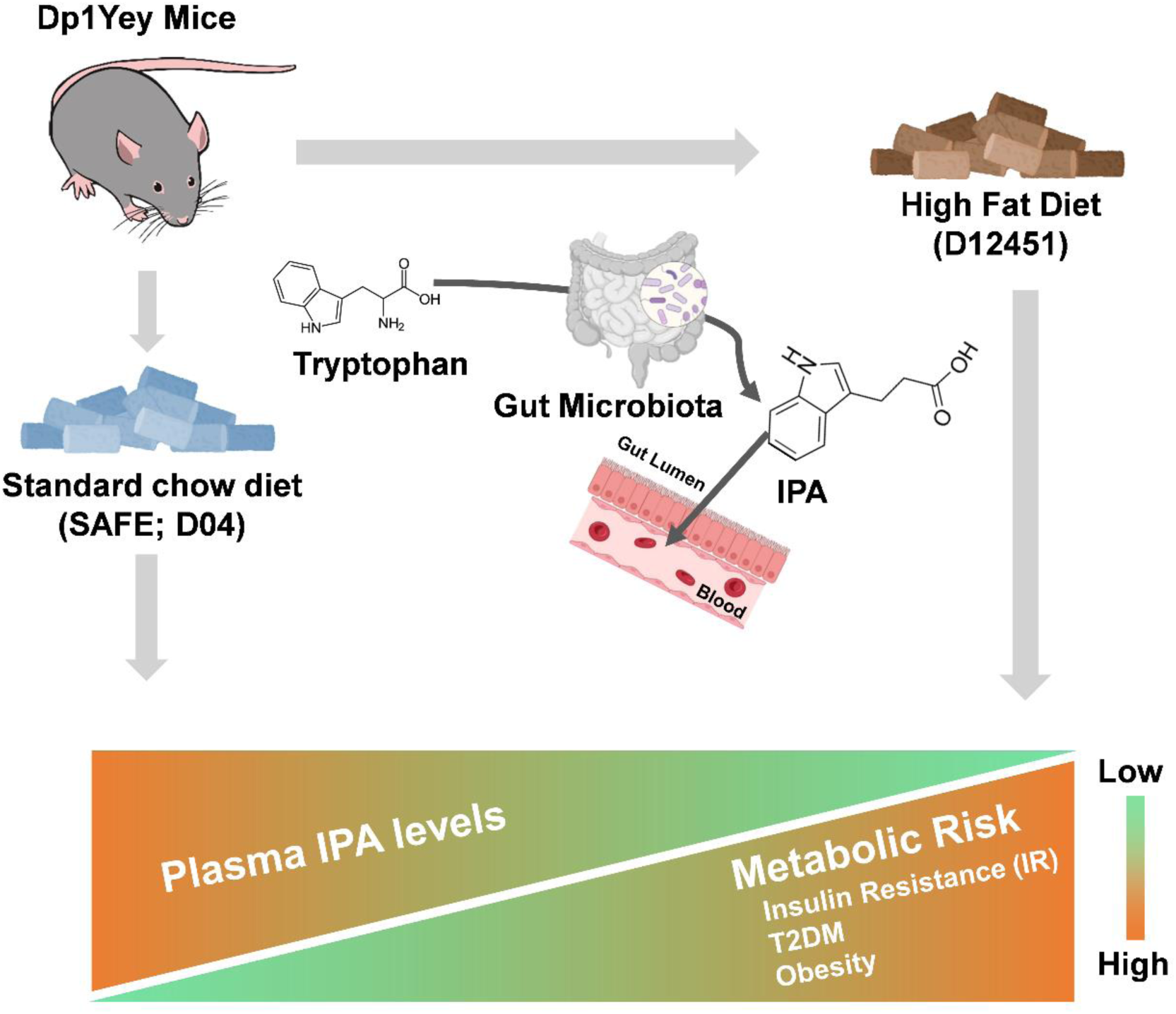
Graphical summary illustrating the role of 3-indolepropionic acid (IPA) in metabolic regulation and its reduction in Dp1Yey mice under high-fat diet. IPA, a gut microbiota-derived metabolite of tryptophan, exerts protective effects by enhancing insulin sensitivity, reducing the risk of type 2 diabetes mellitus (T2DM) and obesity, preserving gut barrier integrity, and mitigating oxidative stress. In Dp1Yey mice, high-fat diet (HFD) feeding results in a specific reduction of circulating IPA levels, indicating disruption of the microbiota–gut–host axis. This decrease may contribute to increased susceptibility to insulin resistance, T2DM, obesity, and broader metabolic dysfunction in the context of segmental trisomy and dietary stress.

## Supplementary Materials

**TableS1**: Metabolites significantly altered by Sex variable, **Table S2**: Metabolic pathways significantly enriched across variables using MetaboAnalyst 5.0, **TableS3**: Metabolites significantly altered by segmental trisomy in the Dp1Yey mouse model, **TableS4**: Metabolites significantly altered by dietary condition, **Table S5**: Metabolic pathways significantly enriched across overlapping variables using MetaboAnalyst 5.0, **Table S6**: Ancova result showing five metabolites that altered due to synergistic effect of Diet*Genotype variable, **Figure S1**: Box plots of significantly altered plasma metabolites stratified by sex, **Figure S2**: Box plots of significantly altered plasma metabolites associated with segmental trisomy, **Figure S3**: Box plots of significantly altered plasma metabolites stratified by dietary condition, **Figure S4**. Plasma IPA levels are selectively reduced in Dp1Yey mice under high-fat diet and show sex-dependent correlations with body weight.

**Figure S1. Box plots of significantly altered plasma metabolites stratified by sex.**

(A) Metabolites elevated in female mice relative to males. (B) Metabolites reduced in females compared to males. Linear models were fitted for each metabolite with Sex as the factor of interest, adjusting for Diet, Genotype and body weight (BW). Type II ANOVA F-tests were performed to extract p-values for the Sex term, followed by BH correction for multiple testing. Significance threshold was set at adjusted *p* < 0.05, and significance levels are indicated as follows: *p* > 0.05 = ns*; p* ≤ 0.05 = *; *p* ≤ 0.01 = **; *p* ≤ 0.001 = ***.

**Figure S2. Box plots of significantly altered plasma metabolites associated with segmental trisomy.** (A) Metabolites elevated in Dp1Yey mice compared to wild type mice (WT). (B) Metabolites reduced in Dp1Yey mice relative to WT. Linear models were fitted for each metabolite with Genotype as the factor of interest, adjusting for Diet, Sex and body weight (BW). Type II ANOVA F-tests were performed to extract p-values for the Genotype term, followed by BH correction for multiple testing. Significance threshold was set at adjusted *p* < 0.05, and significance levels are indicated as follows: *p* > 0.05 = ns*; p* ≤ 0.05 = *; *p* ≤ 0.01 = **; *p* ≤ 0.001 = ***.

**Figure S3. Box plots of significantly altered plasma metabolites stratified by dietary condition.** (A) Metabolites elevated in the HFD group compared to the CTL group. (B) Metabolites reduced in the HFD group compared to the CTL group. Linear models were fitted for each metabolite with Diet as the factor of interest, adjusting for Genotype and Sex. Type II ANOVA F-tests were performed to extract p-values for the Diet term, which were subsequently corrected for multiple testing using the BH method. Adjusted *p*-values < 0.05 were considered statistically significant. Significance levels are indicated as follows: *p* > 0.05 = ns; *p* ≤ 0.05 = *; *p* ≤ 0.01 = **; *p* ≤ 0.001 = ***. Abbreviations: CTL, control; HFD, high-fat diet.

**Figure S4. Plasma IPA levels are selectively reduced in Dp1Yey mice under high-fat diet and show sex-dependent correlations with body weight.** (A) Box plots display log₁₀-transformed plasma IPA peak areas stratified by Genotype (Dp1Yey vs. WT), Sex (Male vs. Female), and Diet (CTL vs. HFD). A three-way analysis of variance revealed a significant main effect of Genotype (*p* < 0.0001) explaining 67.4% of the variance, and significant interactions for Genotype × Diet (*p* = 0.0001) and Genotype × Sex (*p* = 0.026). Tukey’s post hoc comparisons indicated that Dp1Yey-HFD mice exhibited significantly lower plasma IPA levels compared to all other groups (*adjusted p* < 0.0001) particularly in Males. Significance levels are indicated as follows: *p* > 0.05 = ns; *p* ≤ 0.05 = *; *p* ≤ 0.01 = **; *p* ≤ 0.001 = ***; *p* ≤ 0.0001 = ****. (B) Scatter plots of body weight versus plasma IPA levels, stratified by genotype and sex, with regression lines and Spearman correlation coefficients shown above each plot. Significant negative correlations were observed in Dp1Yey males (ρ = −0.678, *p* = 0.0007), WT females (ρ = −0.576, *p* = 0.0026), and WT males (ρ = −0.479, *p* = 0.0099). No significant correlation was detected in Dp1Yey females (ρ = −0.234, *p* = 0.37). Abbreviations: CTL, control; HFD, high-fat diet; WT, wild type.

## Ethics approval

All animal experiments were conducted in accordance with European Union Directive 2010/63/EU (September 22, 2010) and were approved by the ethics committee of the GODS21 project (approval number: 30859).

## Consent for publication and competing interests

All authors have approved the manuscript and consent to its submission and have no conflicts of interest to disclose.

## Data Availability Statement

The original contributions presented in this study are included in the article and Supplementary Material. The raw metabolomics data have been submitted to the MetaboLights repository and are currently under curation (Accession number: MTBLS9532, https://www.ebi.ac.uk/metabolights/reviewercc41a853-8629-4964-9b3a-bd37aaabd5b1). The data will be made publicly available upon completion of the curation process. The raw metagenomics data have been deposited on ENA/Zenodo under the Accession Number doi.10.5281/zenodo.17927869.

## Author Contributions

Conceptualization: MCP, YH; Data curation: PF, FI, GP, YH; Formal analysis: PH, FI, LD, FXL, LL, GP, YH; Funding acquisition: MCP, YH; Investigation: PH, MS, FI, GP, YH, MCP; Methodology: FI, MS, LL, LD; Project administration: MCP, YH; Supervision: MCP, YH; Validation: PH, FI, GP, YH, MCP; Visualization: PH, FXL, LL, GP; Writing—original draft: PH, FI, FXL, MCP, GP, YH; Writing—review and editing: All. All authors have read and agreed to the published version of the manuscript.

## Acknowledgments

We wish to thank the ICAN-Omics platform (https://ihuican.org/en/scientific-platforms/ican-omics/) and the ICM- Data Analysis Core facility (https://dac.institutducerveau.org/). We are also very grateful to the GO-DS21 consortium for constant discussions, particularly with Prof. Pietro Lio.

